# “Degradation of the extracellular matrix is part of the pathology of ulcerative colitis”

**DOI:** 10.1101/412684

**Authors:** Stefan Kirov, Ariella Sasson, Clarence Zhang, Scott Chasalow, Ashok Dongre, Hanno Steen, Allan Stensballe, Vibeke Andersen, Svend Birkelund, Tue Bjerg Bennike

**Affiliations:** Translational Bioinformatics, Bristol Myers Squib, Pennington, NJ, USA; Protein Sciences, Bristol Myers Squibb, Pennington, NJ, USA; Department of Pathology, Harvard Medical School, Boston, MA, USA; Department of Pathology, Boston Children’s Hospital, Boston, MA, USA; Precision Vaccines Program, Boston Children’s Hospital, Boston, MA, USA; Department of Health Science and Technology, Aalborg University, Aalborg, Denmark; Focused Research Unit for Molecular Diagnostic and Clinical Research (MOK), IRS-Center Sonderjylland, Hospital of Southern Jutland, Aabenraa, Denmark; Institute of Molecular Medicine, University of Southern Denmark, Odense, Denmark

**Keywords:** Ulcerative colitis, Proteomics, Extracellular matrix, proteolysis

## Abstract

The scientific value of re-analyzing existing datasets is often proportional to the complexity of the data. Proteomics data are inherently complex and can be analyzed at many levels, including proteins, peptides, and post-translational modifications to verify and/or develop new hypotheses. In this paper, we present our re-analysis of a previously published study comparing colon biopsy samples from ulcerative colitis (UC) patients to non-affected controls. In addition to confirming and reinforcing the original finding of upregulation of neutrophil extracellular traps (NETs), we report novel findings, including that Extracellular Matrix (ECM) degradation and neutrophil maturation are involved in the pathology of UC. The pharmaceutically most relevant differential protein expressions were confirmed using immunohistochemistry as an orthogonal method. As part of this study, we also compared proteomics data to previously published mRNA expression data. These comparisons indicated compensatory regulation at transcription levels of the ECM proteins we identified and open possible new venues for drug discovery.

## Introduction

Ulcerative colitis (UC)is a chronic relapsing inflammatory disorder of the gastrointestinal tract (1–3). The prevalence of UC is increasing worldwide. In Europe, for example, the incidence has reached almost 24 cases per 100,000 person-years (4,5). The disease manifests itself at all ages and it severely affects the quality of life of the patients and their families. In addition, UC has a large impact on society, due to interrupted education, sick leave, and the significant strain it places on the healthcare system.

Both genetics and environment have been found to be factors in the disease etiology. More than 230 genetic loci have been identified as being associated with the inflammatory bowel diseases (IBD) which, in addition to UC, include Crohn’s disease (CD) (6). Identified genes have been found to be involved in the epithelial barrier function and both the innate and adaptive immune response (7). Additionally, a westernized lifestyle and environmental impact is associated with increased risk of IBD, which may be linked to changes in the gut microbiome (8). Accordingly, the prevalence in the West is increasing (9). Cigarette smoking also affects the risk of IBD and gene-smoking interactions have been identified (10).

An early diagnosis and individualized management of the disease are critical in avoiding irreversible damage to the intestinal tract. However, finding good treatments remains a challenge due to the heterogeneous nature of the disease (11–16). Hence, despite the introduction of biological treatments for UC, colectomy still is required in 25-30% of the patients at some point during their lifetime due to the development of colitis refractory to other treatments (17,18). The intestinal mucosa is critical to inflammatory processes and regulates immune responses at the interface between the host and the environment (19–22). We hypothesized that initial damage to the intestinal mucosal barrier results in a misdirected immune response which ultimately leads to the manifestation of IBD (7). An in-depth characterization of the intestinal mucosa via molecular profiling technologies would therefore help to understand the molecular pathophysiology. Using a comparative analysis of the proteome in IBD and healthy colon mucosa, dysregulated molecular analytes and pathways, infiltrating myeloid or lymphocytic markers, chemokines, and inflammatory molecules could be identified. Dysregulated molecular analytes and pathways could be biomarker candidates (prognostic or predictive), diagnostics or even potential targets for drug development.

Recently, we mapped the proteomes of intestinal biopsies from UC patients in remission and gastrointestinally healthy controls, and identified and quantified 5711 different proteins. Our study found an increased abundance of neutrophils and neutrophil extracellular traps (NETs) in the biopsy samples from UC patients (23).

Proteomics data are inherently complex and can be analyzed at many levels. Statistical analysis of proteomics datasets can be confounded by a number of factors. We present here our re-examination of a proteomics dataset comparing UC samples to paired healthy, non-UC controls. In this study we confirmed the findings of the initial publication. In addition, the expanded findings allowed us to hypothesize that the degradation of the colon mucosa extracellular matrix (ECM) may play a critical role in the pathology of UC.

## Materials and Methods

### Analysis reproducibility

All datasets and scripts necessary to reproduce these findings are available from http://pubd.web.bms.com/reproduce.tar.gz.

### Data acquisition

The proteomics raw data were deposited by Bennike et al. (23) to the ProteomeXchange Consortium via the PRIDE ProteomXchange repository with the dataset identifier PXD001608. The study material was composed of colon mucosa biopsies from ten UC patients and ten gastrointestinally healthy controls23s-. These data were retrieved from the PRIDE EBI archive using wget.

### Proteomics search

The sequence database for proteomics searches was assembled from a combination of Ensembl, Refseq and Uniprot databases. In addition, Refseq annotation was used to add fragments unique to mature proteins. We used MaxQuant 1.5.3.8(24) to accomplish all protein identification searches. MaxQuant configurations associated with each search are provided in the Supplementary material section.

The proteinGroups.txt file produced by the MaxQuant search was used for quality control (QC) of the dataset. The QC report was generated using Rstudio(25) running R version 3.3.1 (26). From these initial plots of raw Intensity and LFQ (DP=2), none of the samples was deemed to be of significant concern and thus all samples were included in the subsequent analysis.

### Statistical models

#### Filtering

We removed all proteins where too many data points were missing, using both sample-frequency (66% per group) and donor-frequency (70%) thresholds. Since we had 30 samples per group, a protein that had no readout in 10 or more samples in each group would be considered non-quantifiable and would be eliminated from further analysis. Similarly, proteins considered for further analysis had to be present in at least 7 donors in either group.

#### Imputation

We imputed missing values for quantifiable proteins using the minimum LFQ value in the sample where the missing data point was present. This represents a conservative approach in most cases, as such imputation of values missing at random would likely increase the variability within a group and therefore increase the p-value.

#### Differential expression analysis

We fit a linear mixed-effects regression model of Intensity, with a fixed effect of DiseaseState (UC versus Not) and a random Donor effect, using function lme from the R package “nlme”(27). We controlled for false discovery by applying the Benjamini-Hochberg correction method. Proteins were considered to be significantly differentially regulated at 5% FDR and ±0.5 log2-fold-change. Peptides were considered to be significantly differentially regulated at 5% FDR and ±1 log2-fold-change.

#### Differential modification analysis

To test for the presence of global shifts in modifications (hydroxyproline oxidation, ubiquitination), we averaged the raw intensity from all peptides carrying the specific modification for each donor within the control and UC group. We then applied a two tailed t-test to those averaged values for each group.

#### Microarray data and correlation analysis

Microarray data were retrieved from GEO. Three datasets were reanalyzed: GSE38713(28) (referred to as Plannell), GSE10616(29) (referred to as Kugathasan), and GSE9452(30) (referred to as Olsen). Each dataset was RMA normalized. Only probesets with an expression level more than log2 RMA of 3.5 in at least 10 samples were considered. Fold change values were determined for the control subjects vs the UC patients.

Fold change values for probesets were collapsed to a gene level by taking the median. Pearson pairwise correlations were then calculated for all microarray sets and the proteomics data. The list of proteins with significant disagreement between RNA expression and protein levels was compiled by thresholding for mass spec fold change and microarray median fold change and sorting by the mass spec fold change.

The directionality and downregulation of proteins was stored as a Boolean vector.

#### Pathway analysis

Differentially regulated peptides were converted to protein identifiers using the peptides.txt file. Protein identifiers were converted to EntrezGene identifiers using a combination of conversion techniques according to the source of the originating protein ID (Refseq, Ensembl and Uniprot). We split the lists into up- and down-regulated sublists and examined each list separately with WebGestalt and GSEA. Webgestalt(31) was used to explore GO enrichment and GSEA was used to explore other ontologies and gene lists, such as MSIGDB, Reactome (www.reactome.org) and Pubmed (www.ncbi.nlm.nih.gov/pubmed/).

Both tools allowed lists of genes to be uploaded and to serve as background models and we used all identified proteins to define the background reference.

#### Reproducibility

Most results from this analysis (hits lists, graphs, etc.) can be reproduced on a Linux system with R 3.3.1, perl and several R packages (listed in Supplementary material). Once all input datasets are downloaded along with the scripts, executing reproduce.sh would generate identical results to those we communicate in this paper.

#### Immunohistochemistry

Formalin-fixated colon biopsies (23) were paraffin embedded, cut in 10 µm sections, mounted on Superfrost Plus slides (Thermo Scientific) and deparaffinized. Sections were blocked with 0.1% bovine serum albumin (BSA) in phosphate-buffered saline (PBS) for 1 hour. Thereafter, the sections were incubated for one hour with the primary antibody in 0.1% BSA in PBS using either 2 µg/ml rabbit anti-Protein S100-A9 (Abcam, Cambridge, UK), 2 µg/ml mouse IgG1 anti-Protein S100-A12 (Abcam), 2 µg/ml rabbit anti-myeloperoxidase (ab9535, Abcam), 2 µg/ml isotype control mouse monoclonal IgG1, or 2 µg/ml rabbit IgG. Sections were washed 3 times with PBS. The samples were then incubated for 1 hour in PBS with the secondary antibody using either TRITC conjugated goat anti-rabbit IgG or TRITC conjugated goat anti-mouse IgG (Jackson ImmunoResearch, Bar Harbor, ME). The samples were then diluted 1:200 in PBS with 0.1% BSA. To stain for DNA, the sections were washed in PBS and incubated for 30 minutes with 1 µM ToPro-3 (Thermo Scientific) in PBS. The slides were analyzed with an SP5- confocal microscope (Leica, Wetzlar, Germany) using an HC PL APO 63x/1.40 Oil objective (Leica) and the data was analyzed in ImageJ(32).

## Results

### Quality control

We have established an R-based pipeline to comprehensively evaluate the quality of proteomics data (manuscript in preparation). A full quality report is available in the Supplementary material. We used the quantro R package (33) to evaluate global effects and observed no statistically significant change in protein intensities between the UC and control groups (p=0.11, 10000 simulations). We therefore applied median centering as a global normalization method.

### Protein level analysis

This study aimed to identify proteins differing in abundance on average between the UC and control groups. For this purpose we performed linear regression analysis based on the protein groups generated by MaxQuant.

We identified a similar number of protein groups - 6399 - compared to the original study where 6268 were identified. After rigorous filtering, 5093 proteins were considered to be quantifiable, versus 5711 in the original publication. However, our statistical model allowed us to identify substantially more differentially regulated proteins: 251 versus 49 in the previous study. This difference reflects two methodological differences: different filtering and imputation strategies, and the linear mixed effects model we used that allowed us to take advantage of the technical replicates, which had been averaged in the original study. Including a random donor effect to allow for intra-donor correlations between technical replicates allowed the assessment of technical variability and increased statistical power of hypothesis tests.

As with the original study we were able to establish statistically significant overlap with NETs (hypergeometric test, p<0.001), most significantly with NET granules(34). We were able to confirm the presence of a neutrophil immune cell signature also based on MSIGDB(35).

Of the differentially regulated proteins, 90 had increased and 161 had decreased abundance in the UC samples. However, 9 of the top 10 proteins with the largest absolute expression difference (Table 2) were upregulated, signifying the larger effects observed in diseased tissue. The only downregulated protein in the top 10 is elastin, an indication of changes in the extracellular matrix. This observation is in line with previous research implicating bacterial or host proteases in the degradation of the ECM in IBD(36,37).

**Table 1:** Number of participants in the three microarray studies

| Study | Controls | Ulcerative colitis |
| --- | --- | --- |
| Olsen(30) | 5 | 21 |
| Planell(28) | 13 | 30 |
| Kugathasan(29) | 16 | 10 |

**Table 2:** The top 10 differentially regulated proteins when comparing the UC and control groups. The list is filtered by FDR adjusted p-value (p<0.05) and sorted by effect size (Log2 ratio).

| **Protein Acc.** | **p-value** | **FDR** | **Entrez gene** | **Log2 Ratio** | **Description** | **Gene** | **NETs protein** |
| --- | --- | --- | --- | --- | --- | --- | --- |
| NP_002334 | 8.57E-12 | 4.37E-08 | 4057 | 7.45 | Lactotransferrin | LTF | Yes |
| XP_011523124 | 2.02E-13 | 1.03E-09 | 4353 | 6.76 | Myeloperoxidase | MPO | Yes |
| NP_004985 | 1.87E-12 | 9.52E-09 | 4318 | 6.06 | Matrix metalloproteinase 9 | MMP9 | Yes |
| XP_011545190 | 2.93E-07 | 1.49E-03 | 1991 | 5.02 | elastase, neutrophil expressed | ELA2 | Yes |
| NP_001691 | 1.28E-08 | 6.50E-05 | 566 | 4.92 | azurocidin 1 | AZU1 | Yes |
| XP_011514177 | 5.16E-13 | 2.63E-09 | 2006 | -4.88 | Elastin | ELN | No |
| NP_002956 | 2.13E-09 | 1.09E-05 | 6280 | 4.67 | S100 calcium binding protein A9 | S100A9 | Yes |
| NP_000623 | 2.02E-10 | 1.03E-06 | 3684 | 4.53 | integrin subunit alpha M | ITGAM | No |
| NP_002768 | 6.37E-07 | 3.24E-03 | 5657 | 4.49 | Proteinase 3 | PRTN3 | Yes |
| XP_005260579 | 8.75E-10 | 4.46E-06 | 671 | 4.45 | Bactericidal/permeability-increasing protein | BPI | No |

We compared our results to previous research of differential gene expression in UC. For that purpose we selected 3 publicly available transcriptional profiling datasets(28–30). We extracted the lists of differentially regulated genes from the microarray data and then calculated the log2 ratio for all sets, including the present proteome dataset. All further comparisons were based on this metric. Merging between datasets was done using a translation table available in the Supplementary material. We were able to match 219 of the 251 hits in all datasets. Overall the correlation between the transcriptional profiling studies is high. In contrast, the correlation to our proteomics dataset was moderate with R<=0.38 (Figure 1). Unlike mRNA-protein level correlation, for which we anticipated low correlation based on established research (38), we expected differentially expressed proteins to exhibit greater correlation when fold changes were used for comparison (39).

**Figure 1:**
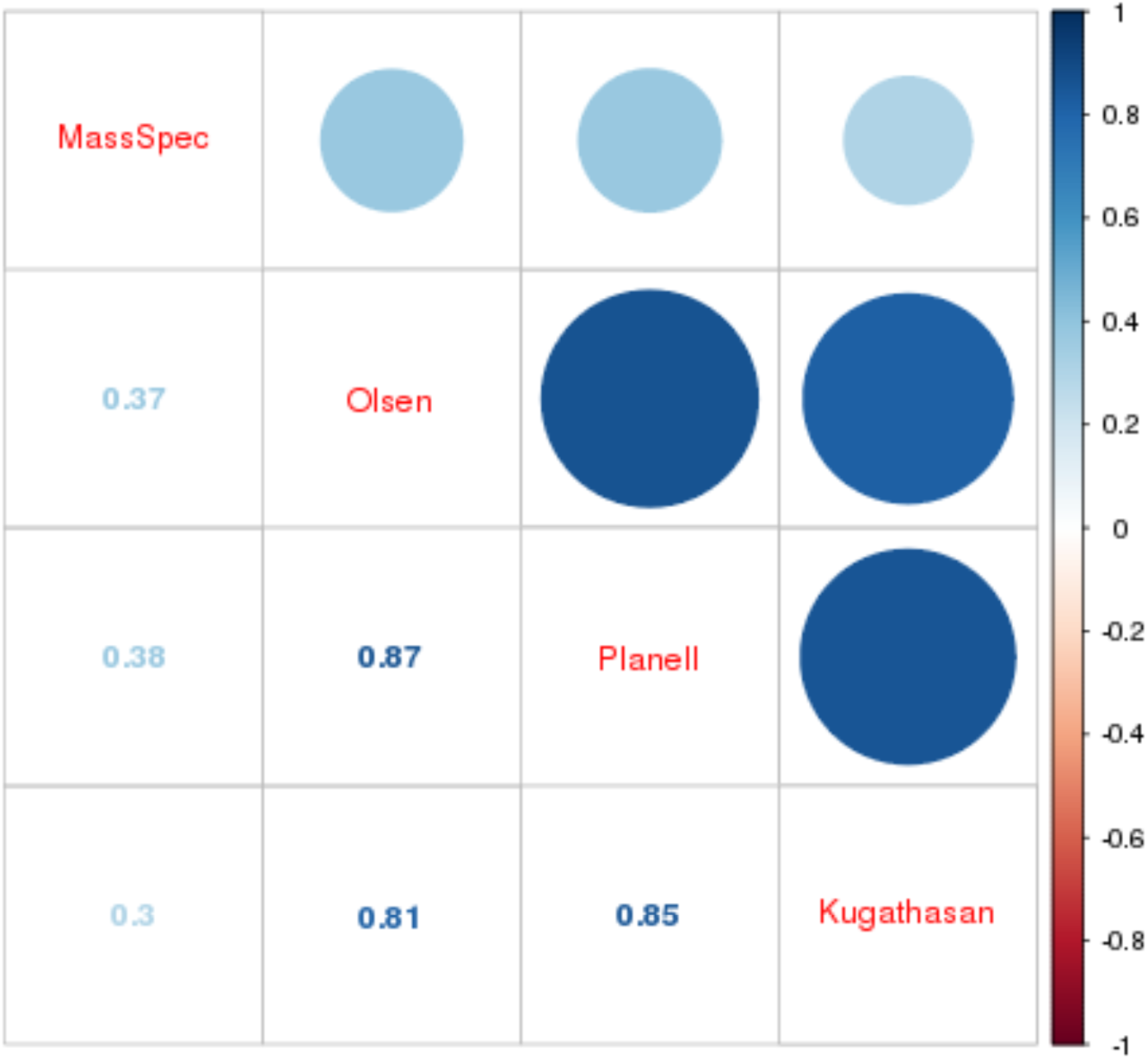
The correlation matrix based on Pearson correlation of log2 ratios.

Surprisingly, genes that were highly expressed in neutrophils and are considered granulocyte markers showed different direction changes in the proteomics data and even between different transcriptional studies. Myeloperoxidase (MPO), for example, is well studied and known to be highly induced in active UC(40). The reverse observation may be the result of a suboptimal design of microarray probesets. The lack of high correlation may be due to post-translational events, differences in general technologies, or sample selection. Another possible explanation would be that transcription is halted when secretory granules are full, resulting in high protein levels, but low transcript abundance. There also were significant differences between the transcriptional and protein fold changes when examining the downregulated proteins (Supplementary material). A substantial number (49) of the 149 downregulated proteins were either upregulated (log2 ratio>0.5, n=16) or unchanged with log2 ratio between 0 and 0.5 (n=33). Most of these proteins (Table 3) are part of the ECM. Anecdotal reports of similar phenomena have been recorded in the past. For example, collagen protein is known to be lost due to degradation in models of IBD, and PGP fragments are subsequently increased (41). This degradation of collagen could trigger compensation at a transcriptional regulation level and would explain the upregulation seen in the microarray studies (28–30) we examined.

**Table 3:**
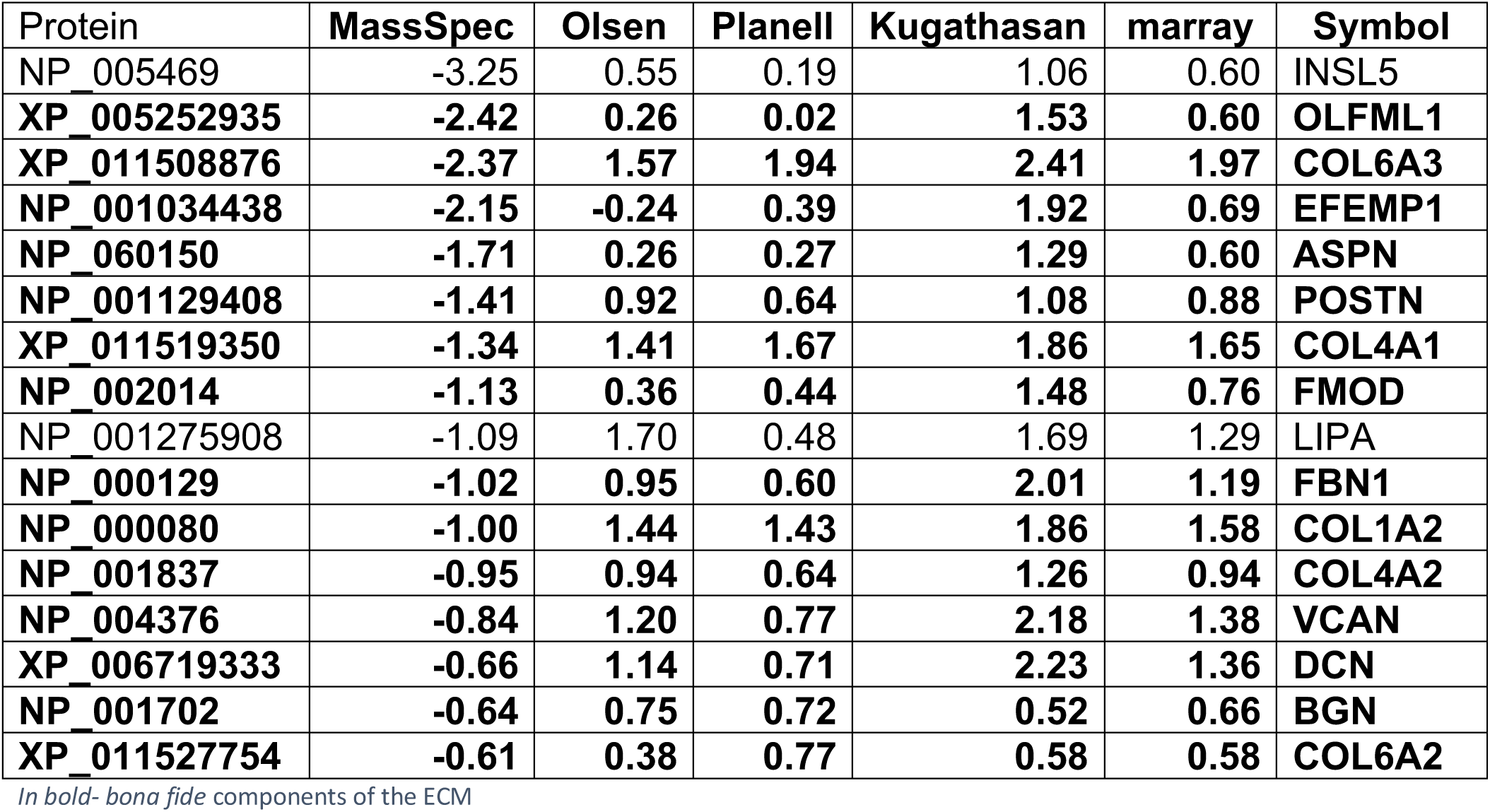
Top 10 of the proteins where significant disagreement between protein log2 ratios and transcriptional profiling ratios were observed, ranked by mass spec log2 fold change

### Chromogranin A

In addition to the protein level analysis, we also looked at differential expression at the peptide level (Supplementary material). This analysis may occasionally highlight differences in protein processing and secretion. We decided to look specifically at chromogranin A, as the log2 ratio for the identified peptides originating from this protein showed substantial dispersion.

Chromogranin A (CHGA) is a protein that is post-translationally processed to multiple hormones and functional peptides(42,43). While chromogranin A was one of the differentially regulated proteins, it was not a top hit. CHGA ranked in the top 28.17% of all hits (71/252) in terms of fold change (log2 FC=-1.77, FDR=0.0015). At the peptide level, differential regulation was more pronounced with fold changes ranging from −3.7 to −5.7, and the top peptide ranked at the top 6.54% of all hits (42/642). The reason for this discrepancy is that some of the peptides were significantly downregulated in UC (Figure 2), while others were only marginally higher in controls. For all peptides, the characteristics and fold change differences between tissues from healthy controls and UC patients are given in the Supplementary material.

**Fig 2:**
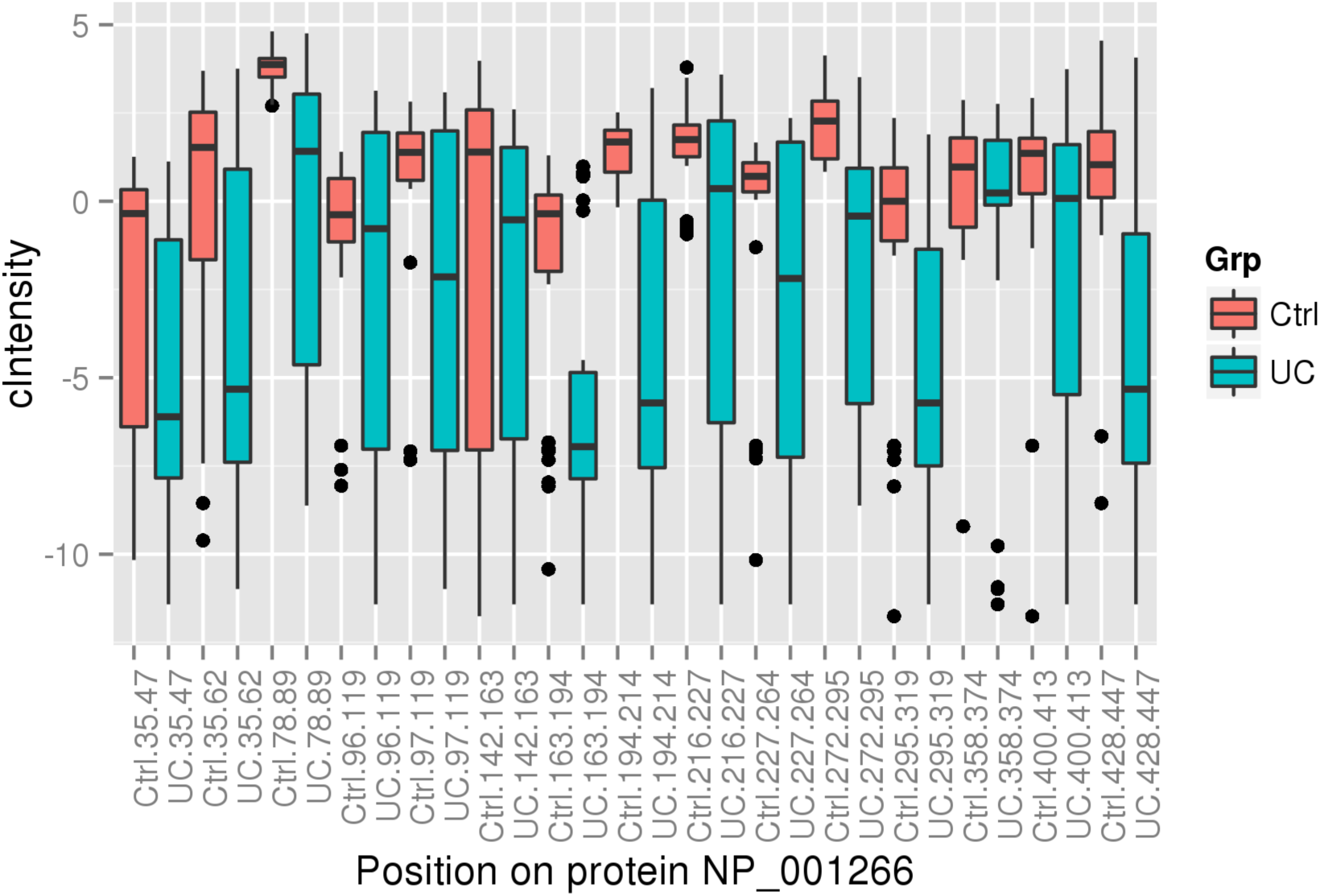
Mean normalized intensity of chromogranin A peptides in UC and controls according to the position of the individual peptides in the full protein sequence

Identification of CHGA sub-regions as some of the most significant peptide hits (Supplementary material) allowed us to describe EA-92, chomofungin, serpinin, and pancreastatin as potential biomarkers and perhaps targets in the context of UC. In addition, we can now speculate on the involvement of EA-92 in inflammation.

Significant changes occured in four peptide regions. These regions overlap with the biologically active peptides serpinin, vasostatin and chromofungin that are known to have anti-microbial activity(44,45). As we did not intend to explore bioactive peptide expression, it is hard to define exactly which functional regions are differentially regulated. We can speculate, for example, that the CHGA downregulated peptide probably originates from chromofungin, since changes in other vasostatin I and vasostatin II peptides were not statistically significant. We also observed significant changes in EA-92, but some peptides originating from the same region were not consistent with this change. It is possible that EA-92 is further processed into a shorter mature product.

### Decreased Hydroxyproline in Ulcerative Colitis

Collagen is a central ECM protein and degradation of it has been associated with IBD(41). Hydroxyproline modifications correlate well with collagen content, collagen protein abundances, and the degree of fibrosis(46) which can be found in >30% of CD patients and about 5% of UC patients(47,48). We therefore decided to include a quantitative analysis of the hydroxyproline PTM content. In addition, ubiquitination has been implicated in IBD and was for that reason also included in the analysis(49).

We did not observe differential regulation of any of the individual amino acid sites, either for hydroxyproline or ubiquitinated PTMs. Additionally, we did not observe a statistically significant difference between total or peptide specific ubiquitination (p=0.146). However, the overall abundance of peptides with a hydroxyproline modification was significantly lower in the UC colon tissue compared to control (Figure 3), which is consistent with the reduced abundance of collagen proteins determined by proteomics and previously published research of collagen degradation(50).

**Fig 3:**
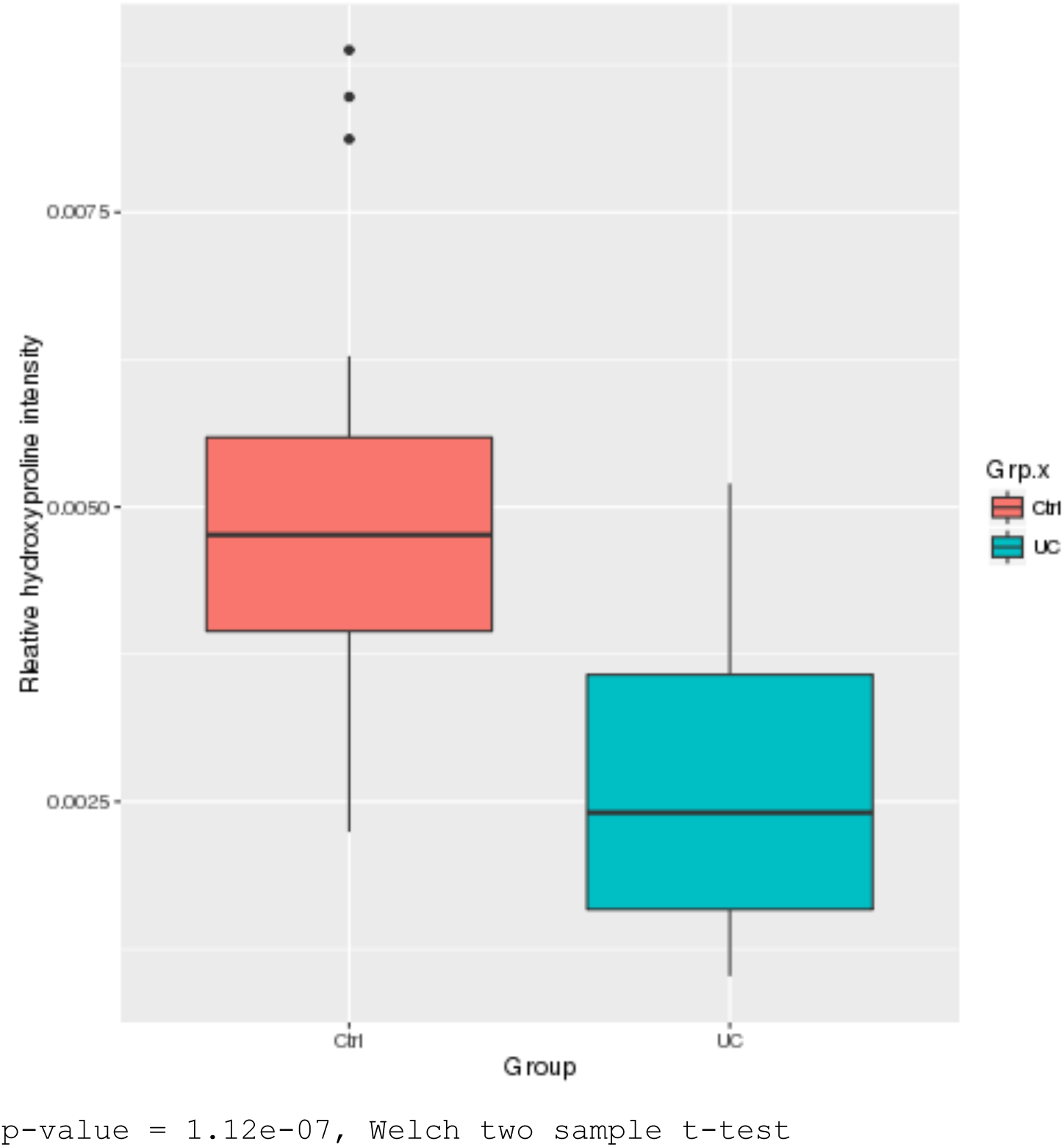
Overall change in hydroxyproline modified peptide intensity

### Pathway analysis

We explored the presence of over-represented categories from several different ontologies using lists of upregulated and downregulated features at the peptide and protein levels. When using the GO Biological process ontology and comparing to our protein-level list (Supplementary material), most categories were related to the immune system. The most significant category we observed was “Response to bacteria”. We also observed the enrichment of leukocyte migration, which is a hallmark feature of IBD(51). At the same time, downregulated proteins significantly overlapped with extracellular structure organization as well as mitochondrial respiratory complex assembly and glycosylation (Supplementary material). While mitochondrial changes have been previously implicated in the pathogenesis of IBD(52), the association of ECM categories with downregulated proteins was largely unexpected. This observation was consistent with the decreased hydroxyproline levels and the data we obtained by comparing transcriptional profiling data with the list of differentially regulated proteins.

Next, we explored additional ontologies and gene lists, including MSigDB and Pubmed (Supplementary material). Most lists we identified as significantly overlapping with our list of upregulated proteins were related to immune function, and further reinforce the neutrophil association. One category also of interest is Reactome’s “Degradation of extracellular matrix”, which is consistent with decreased hydroxyproline levels and collagen protein, supporting our hypothesis. In addition, it appears that lists associated with cell line proliferation (stem cells, cancer) overlap significantly with our downregulated list of proteins.

In the GO Component ontology,only one category was enriched in the list of upregulated peptides: “Blood microparticles” (Supplementary material). Blood microparticles (MPs) have been associated previously with inflammation and infection, including Crohn’s disease(53). Secreted MPs or secreted Microvesicles (sEV) are a diverse range of lipid vesicles containing a cargo of protein and RNA/DNA originating from a range of cell types including neutrofils (54).

### IHC confirmation

Colon biopsy sections from 4 UC patients and 2 healthy controls were stained for the proteins S100-A9 and S100-A12, or for myeloperoxidase, all of which are present in granulocytes. In Figure 4, results from one patient with ulcerative colitis and one healthy control are shown. No reaction with antibodies to S100-A9 and S100-A12 was observed in sections of the healthy control (Figure 4, A and D) or in immunoglobulin controls (Figure 4, F and I), showing that granulocytes are not present in healthy intestinal mucosa (mucous epithelium and lamina propria). A strong staining was observed for S100-A9 in the UC patients, and, in addition, more cells were observed in lamina propria than in controls. (Figure 4, B). By enlarging the image, it can be seen that cells stained for S100-A 9 are polymorphonuclear (Segmented nuclei), in agreement with granulocytes, and S100-A9 is located in the cytoplasm of the cells (Figure 4, C). It can also be seen that the S100-A9 contained in polymorphonuclear cells are located between the mucosal epithelial cells (Figure 4, B and C). S100-A12 are also present in the cytoplasm of polymorphonuclear cells in the UC patient (Figure 4). Myeloperoxidase shows a similar staining pattern as S100-A9 and S100-A12, in agreement with its localization in granulocytes.

**Figure 4:**
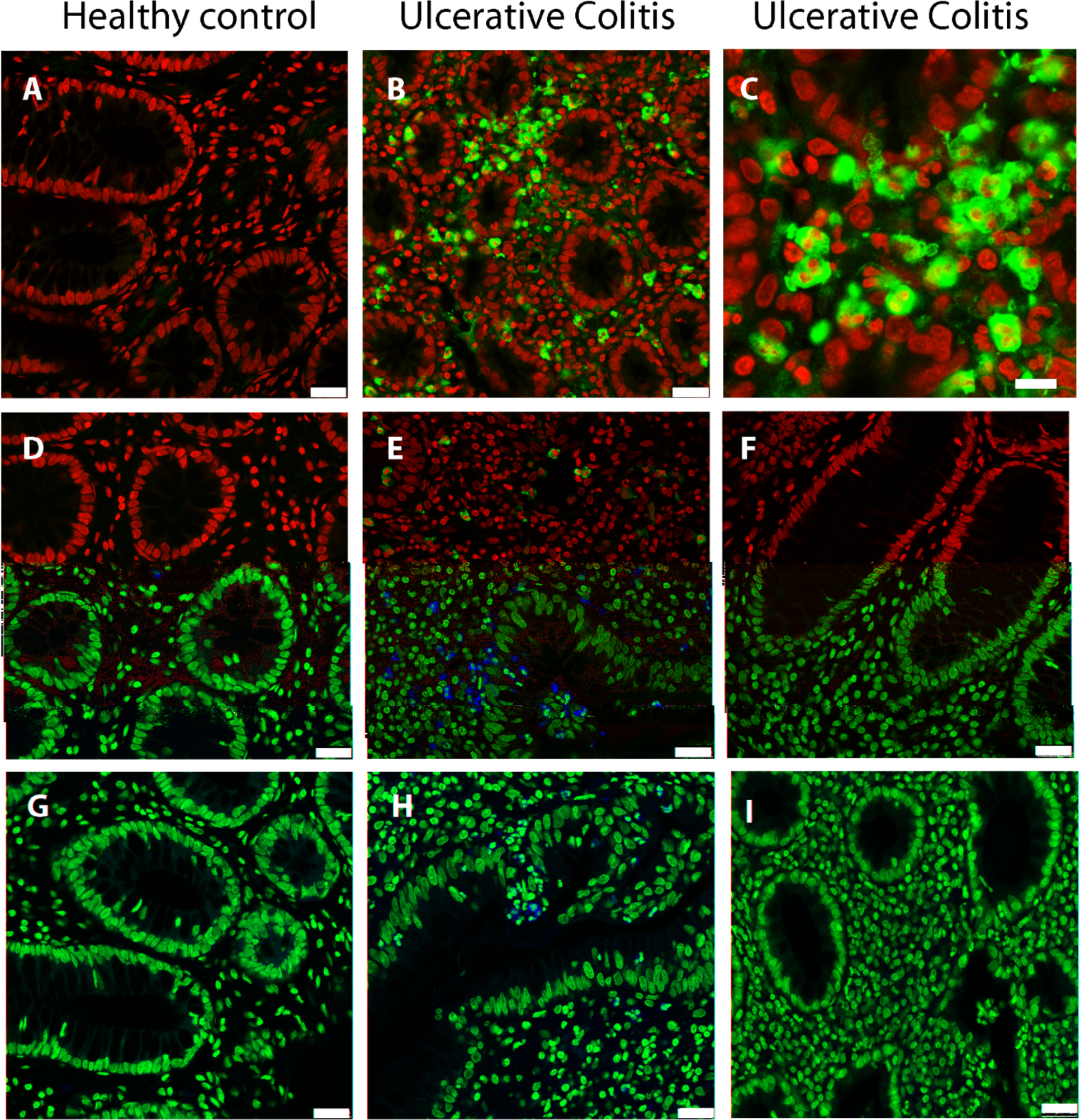
Confocal microscopy of colon biopsies from a healthy control and an ulcerative colitis patient. A) Healthy control, B) and C) Ulcerative colitis patient stained with anti-protein S100-A9. Healthy control D) and E) Ulcerative colitis patient stained with anti-protein S100-A12. F) Isotype control with mouse IgG1 monoclonal antibody. G) Healthy control stained with anti-myeloperoxidase and H) Ulcerative colitis patient. I) IgG control of ulcerative colitis patient section stained with normal rabbit IgG. Nuclei are stained red with To-Pro3. The bar indicates 25 µm in the picture A, B, D-I and 10 µm in C.

## Discussion

One of the most unexpected results we report here, supported by several separate findings, is the presence of significantly lower levels of multiple ECM proteins in intestinal biopsies from UC patients. Such downregulation can occur as a result of either modulation of transcriptional and translational activity or degradation and displacement (with the second explanation being more likely based on microscopy findings). The notion that degradation drives the changes we observe is further supported by the increased detection of ECM matrix fragments, reported in the serum of UC patients vs controls or Crohn’s disease patients (55,56).

Fibrosis is a well described complication of IBD (57), especially in CD(47,48). Therefore, we expected to observe upregulation of collagen proteins and hydroxyproline-containing peptides. Indeed, increases in collagen transcripts were found with all 3 previously published transcriptomics datasets (28,30,58). However, among current proteomics datasets several collagen proteins were found to have lower abundances in the UC group. The discovery of decreased collagen protein abundance in UC was supported by a lower abundance of hydroxyproline-containing peptides, and was in agreement with previous reports of UC colon mucosa(59). The UC patients in our proteomics study were all medicated and no visible inflammation could be observed on the surface of the colon from which the biopsies were taken. Additionally, the histology analysis of the colon biopsies found that the crypt architecture was preserved and without distinct fibrosis, which is in agreement with the proteomics findings. However, an increased presence of neutrophils, NETs, and proinflammatory proteins could be detected. Taken together, the findings indicate the presence of a low-grade inflammation in the tissue, which could trigger the degradation of collagen.

We speculate that these seemingly contradictory findings between the collagen protein abundances and the transcript levels are two sides of the same complex process. It is possible that an active process caused by or causing the degradation of the ECM would lead to a compensatory yet insufficient transcriptional response. It would be interesting to learn the turnover of ECM proteins in the active sites of the disease. Increased rates would support our hypothesis, and increased ECM protein turnover has already been observed in the bleomycin model of pulmonary fibrosis(60).

Consistent with ECM degradation, we detected increased abundances of ECM degrading enzymes, including myeloblastin, neutrophil collagenase, MMP9, and MMP10. However, our data somewhat surprisingly cannot confirm that MMP9 and MMP10 upregulation is the cause for the ECM protein degradation as we could not detect increased abundance of the activated MMP fragments, suggesting a lack of MMP activation. This observation requires further validation as we are basing this hypothesis on known MMP substrates which may not represent the full diversity, thus potentially skewing the results. In addition, it is possible the current proteomics workflow may not be able to assess the presence of MMP fragments fully. However, the increase in MMP proteins is consistent with the reduced abundance of collagen proteins and hydroxyproline, and by extension the reduction in ECM protein content, thus adding credibility to the potential of ECM degradation as a major event in IBD pathology. We can further speculate that ECM degradation is an initial trigger for both neutrophil infiltration and ECM transcriptional activation which may overcompensate and eventually result in fibrosis. Purely speculatively, ECM degradation could be triggered partially by leaky epithelial tight junctions in intestinal inflammation. Tight junctions are multi-protein complexes that form a seal between adjacent epithelial cells, and thus act as barriers that normally regulate the transport of macromolecules. UC genetic studies have identified susceptibility loci related to defects of the epithelial barrier (61,62), and IBD patients often demonstrate a loss of tight junction functions. While not sufficient to cause IBD, the loss of the tight junction function causes an immune activation(63). Accordingly, epithelial leaks can be detected early in UC(64), and it has been suggested that these barrier defects allow bacterial antigens to enter the mucosa from the lumen, causing inflammation of the mucosa which could lead to ECM degradation(65).

Degradation of ECM proteins could potentially have an impact on IBD pathology. It has been reported that proteolytic fragments (PGP) of ECM proteins can serve as neutrophil chemoattractants through CXCR1(66). PGP has also been identified as a potential biomarker in COPD, another disease where inflammation and neutrophil infiltration play a major role(67).

In conclusion, emerging evidence of protein degradation and peptidase involvement in the pathology of UC may open new opportunities for biomarker discovery and novel therapeutic options. We also observed another interesting example of disconnect between protein and mRNA regulation of Chromogranin A (CHGA). CHGA is a great example of a situation where the mRNA and its role in the generation of multiple hormones was characterized decades ago(42,68,69), yet we continue to find novel proteoforms originating from this locus(44). Chromogranin A upregulated peptides also diverge significantly in terms of differential regulation, potentially due to the different regulation and/or secretion of the many proteoforms that are encoded by this locus. And while chromogranin A itself is differentially regulated, specific regions of the protein exhibited much larger differences between UC patients and the control population.

An elevated level of NETs is closely associated with general inflammation by the release of a extracellular lattice, composed of DNA associated with proteins citrullinated by protein-arginine deiminase 4 (PAD4) from neutrophils (70). The NETosis or NETlysis processes are associated with a range of autoimmune diseases or triggered by inflammatory activation including UC and reumatroid arthritis (23,71). Berthelot et. al. have found that NETosis increases the possibility of association between autoantigens and infectious antigens in mucosal biofilms, impairing the clearance of pathogens and possibly triggering autoimmune reactions such as autoantibody formation (ANCAs). These autoantibodies against NET components have been suggested as possible explanations for the breakdown of tolerance to NET autoantigens, such as hypercitrullination. Of the NETs granule proteins identified in our study, LTF, MPO, and PRTN3 have been associated with autoantibody formation in relation to IBD and ECM-related processes (72). The release of active PAD2 and PAD4 isoforms into synovial fluid by neutrophil cell death is a plausible explanation for the generation of extracellular autoantigens in RA (73). The inhibition of PADs by either small molecule drugs or biologics has been implicated in treatment strategies for autoimmune diseases such as RA but no evidence for such potential has been provided so far for IBD (74).

One could argue that proteomics data are more complicated than other OMICs data. RNASeq- and DNASeq-data have a single output: variants and/or transcript expression. In contrast, proteomics data can be analyzed in different contexts-expression levels as well as post-translational modifications, and variations. In addition, we have a much better understanding of splicing isoforms than of proteoforms. All this presents a significant challenge to the community. Therefore, in this work, we re-investigated an existing proteomics dataset, and confirmed and extended the original findings. Some of the differences in the data analysis method included variations in the proteomics identification, filtering, and imputation procedures. However, the largest contributor to the new observations and results can be ascribed to the different statistical test that was applied. In the original dataset, statistically significant proteins were identified mainly using t-tests. In comparison, in our reanalysis, we used tests based on linear mixed effects models, which allowed us to include the replicates in the statistical analysis rather than combining them, thereby increasing the number of significant proteins identified substantially. Combining these findings with an existing RNAseq-dataset, we were able to draw novel conclusions, including ECM degradation in UC. In addition to the increased understanding of the pathophysiology of UC, our work demonstrates the significant value in sharing raw data with the community through resources such as PRIDE to enable such cross-OMIC analyses.

